# Multiple steps mediate ventricular layer attrition to form the adult mouse spinal cord central canal

**DOI:** 10.1101/676767

**Authors:** Marco A. Cañizares, Aida Rodrigo Albors, Gail Singer, Nicolle Suttie, Metka Gorkic, Paul Felts, Kate G. Storey

## Abstract

The ventricular layer of the spinal cord is remodelled during embryonic development and ultimately forms the adult central canal, which retains neural stem cell potential. This anatomical transformation involves the process of dorsal collapse, however, accompanying changes in tissue organization and cell behaviour as well as the origin of cells contributing to the adult central canal are not well understood. Here we describe sequential localised cell rearrangements which contribute to the gradual attrition of the spinal cord ventricular layer during development. This includes local breakdown of the pseudostratified organisation of the dorsal ventricular layer prefiguring dorsal collapse and evidence for a new phenomenon, ventral dissociation, during which the ventral-most floor plate cells separate from a subset that are retained in the central canal. Using cell proliferation markers and cell-cycle reporter mice, we further show that following dorsal collapse, ventricular layer attrition involves an overall reduction in cell proliferation, characterised by an intriguing increase in the percentage of cells in G1/S. In contrast, programmed cell death does not contribute to ventricular layer remodelling. By analysing transcript and protein expression patterns associated with key signalling pathways, we provide evidence for a gradual decline in ventral sonic hedgehog activity and an accompanying ventral expansion of initial dorsal bone morphogenetic protein signalling, which comes to dominate the forming central canal. This study identifies multiple steps that contribute to spinal cord ventricular layer attrition and adds to increasing evidence for the heterogenous origin of the adult spinal cord central canal, which includes cells from the floor plate and the roof plate as well as ventral progenitor domain.

## Introduction

The ependymal cells that form the central canal in the adult mammalian spinal cord constitute a largely quiescent stem cell niche (Adrian and Walker, 1962, Kraus-Ruppert et al., 1975, Alfaro-Cervello et al., 2012, Sabourin et al., 2009). These cells can be induced to re-enter the cell cycle in response to extrinsic stimuli, including mechanosensory stimulation (Shechter et al., 2011), physical exercise (Krityakiarana et al., 2010), inflammation (Chi et al., 2006, Danilov et al., 2006) and injury (Adrian and Walker, 1962, Frisen et al., 1995, Barnabe-Heider et al., 2010, Johansson et al., 1999, Li et al., 2016, Li et al., 2018, Meletis et al., 2008). Moreover, most of these stimuli appear to promote the generation of new neurons and glial cells, consistent with *in vitro* studies of the differentiation potential of the spinal cord ependymal cell population (Weiss et al., 1996, Johansson et al., 1999, Li et al., 2016, Meletis et al., 2008, Sabourin et al., 2009). However, following spinal cord injury, central canal cells proliferate and migrate to the lesion site, but here differentiate into only glia (Barnabe-Heider et al., 2010, Li et al., 2016, Li et al., 2018, Meletis et al., 2008, Martens et al., 2002). These cells then contribute to scar tissue, many becoming astrocytes which reduce inflammation, but chronically inhibit axonal re-growth (Warren et al., 2018), while others differentiate into oligodendrocytes, which can promote survival of nearby neurons and help to maintain the integrity of the injured spinal cord (Sabelstrom et al., 2013). Together, these findings indicate that changes in environment determine the behaviour and differentiation of spinal cord ependymal cells. Importantly, this is a heterogenous cell population and the precise identity of cells with neural stem cell abilities has yet to be determined. This activity of spinal cord ependymal cells is also distinct from that of ependymal cells lining the brain ventricles, where instead the neural stem cells constitute a distinct sub-ependymal cell population (Mirzadeh et al., 2008, Shah et al., 2018, Lim and Alvarez-Buylla, 2016). In the healthy animal, adult spinal cord ependymal cells carry out specialised functions, including homeostatic regulation of cerebrospinal fluid (CSF) composition and acting as a barrier between CSF and the spinal cord parenchyma (reviewed in (Del Bigio, 1995, Bruni, 1998)). However, despite these significant roles in the injured and healthy spinal cord, little is known about how spinal cord ependymal cells arise and how the central canal is formed during development.

One aspect of central canal formation involves attrition of the progenitor cell population that constitutes the ventricular layer of the embryonic spinal cord (Fu et al., 2003, Shibata et al., 1997, Yu et al., 2013). This remodelling process includes a striking morphological phenomenon known as dorsal collapse, which mediates a pronounced reduction of the dorsal ventricular layer in a range of mammals (Barnes, 1883, Bohme, 1988, Elmonem et al., 2007, Sevc et al., 2009, Sturrock, 1981). However, the changes in cell behaviour that underlie this critical event are poorly understood. In contrast, the earlier dorso-ventral subdivision of the developing spinal cord has been well-characterised. This involves signals emanating from the roof plate located at the dorsal midline (including bone morphogenetic protein (BMP) and Wnt) and the floor plate at the ventral midline (Sonic hedgehog, Shh), which act in opposition to specify distinct neural progenitor cell populations along the dorso-ventral axis (Jessell, 2000, Le Dreau and Marti, 2012, Ulloa and Briscoe, 2007). This involves regulation of homeodomain and other transcription factors, which act in combination to define neuronal subtype specific progenitors (Lee and Pfaff, 2001). Key transcription factors include *Pax6*, which is initially expressed broadly and is involved in motor neuron progenitor specification, and *Nkx6-1*, which distinguishes V2 interneuron progenitors. Importantly, the continued expression of *Pax6* and *Nkx6-1* in the adult central canal has led to the notion that ependymal cells derive from this earlier population of ventral neural progenitors (Fu et al., 2003, Yu et al., 2013). It is apparent that this ventral region of the ventricular layer is also reduced over time and this may be associated with the switch from neurogenesis to gliogenesis between E11.5-12.5 and, ultimately, the migration of glial cells out of this layer (Deneen et al., 2006, Stolt et al., 2003) reviewed in (Laug et al., 2018). As the emerging central canal becomes separated from the most dorsal and ventral regions of the spinal cord, its formation may additionally involve the remodelling of these specialised cell populations. Indeed, dorsal collapse coincides with elongation of processes from nestin-expressing cells from the roof plate, which ultimately integrate into the adult central canal in mammals (Bohme, 1988, Sevc et al., 2009, Xing et al., 2018, Shinozuka et al., 2019, Ghazale et al., 2019) and fish (Kondrychyn et al., 2013). It is also possible that a similar ventral reorganisation takes place and that this may account for the apparent inclusion of some floor plate cells in the adult central canal (Khazanov et al., 2017).

Here, we describe sequential cell rearrangements associated with the attrition of the ventricular layer as the spinal cord matures during mid to late mouse embryogenesis. We determine whether changing patterns of cell proliferation and/or programmed cell death may contribute to this process. We further evaluate the contribution of floor plate cells to the forming central canal, assessing this in the context of changing expression patterns of genes and proteins associated with key dorsal and ventral signalling pathways.

## Methods

### Animals

Mice were either wild type CD1 or C57BL/6J strains (Charles River). Embryos and spinal cords from Fucci2a transgenic mice (Mort et al., 2014) were provided by Professor Andrew Jackson (MRC HGU, University of Edinburgh) and embryos and spinal cords from Shh-GFP (Chamberlain et al., 2008) and GBS-GFP (Balaskas et al., 2012) transgenic mice were provided by Dr James Briscoe (Francis Crick Institute). For timed matings, the morning of the plug was considered E0.5. All animal procedures were approved by the UK Government Home Office and in accordance with European Community Guidelines (directive 86/609/EEC) under project licence numbers 6004454.

### Immunofluorescence

From embryos, the inter-limb (thoracic) spinal cord tissue was dissected in ice-cold PBS, fixed for 2 hours at 4°C in 4% paraformaldehyde (PFA), rinsed several times in PBS and cryopreserved in 30% sucrose/PBS at 4°C overnight. This tissue was embedded in 1.5% LB agar/5% sucrose, again cryopreserved in 30% sucrose/PBS overnight, frozen on dry ice, and either cryosectioned at 20 μm sections or stored at −20 °C for later use. To obtain adult spinal cords, 10 to 24-week old mice were deeply anaesthetised with an overdose of pentobarbital (Euthatal) or isoflurane and then transcardially perfused with ice-cold PBS followed by 4% PFA. Spinal cords were quickly dissected out and post-fixed in 4% PFA for 2 hours at 4°C and processed as above. Standard procedures were used for immune fluorescence, with the following exceptions. To detect phosphorylated SMAD1/5, all solutions were supplemented with PhospoStop phosphatase inhibitor (Roche, Cat. # 4906837001) and sections were dehydrated with methanol to improve permeability; to detect pH3 sections were exposed to citrate buffer (pH 6) at 95°C for 20 min. In all cases, sections were placed in blocking buffer (2-10% heat-inactivated donkey or goat serum and 0.3-1% Triton X-100 in PBS) for 30 minutes at room temperature and incubated at 4°C overnight with primary antibodies diluted in blocking buffer. Lists of primary and secondary antibodies used are provided in Table S1. Nuclear counterstaining was achieved with DAPI diluted in PBS (1:1000) and incubated for 5 minutes at RT. Slides were mounted with ProLong® Gold Antifade Mountant (Thermo Fisher Scientific, P36930) and images were captured on a DeltaVision or Leica TCS SP8 confocal microscope system.

### RNA *in situ* hybridisation

Inter-limb spinal cord tissue was dissected, fixed and processed as described above, but cryosectioned at 50 μm. Adult mouse spinal cords (24 to 40-weeks old) were also prepared as above but post-fixed overnight in 4% PFA at 4°C. Plasmids, restriction endonucleases and RNA polymerases used for the generation of digoxigenin-labelled probes (Roche Applied Science) are provided in Table S2. Probes were diluted 1:100 in hybridisation buffer (50% deionised formamide, 5X SSC, 0.08% Tween 20, 50 μg/mL heparin, 1 mg/mL t-RNA, 5 mM EDTA, 0.1% CHAPS, 0.02 g/mL Boehringer blocking reagent) and slides incubated at 65°C in a chamber humidified overnight. Sections were washed in post-hybridisation buffer (1X SSC, 50% formamide) at 65°C for 15 minutes three times, followed by a 1:1 post-hybridisation buffer/TBS-0.1% Triton X-100 (post-hyb) wash at 65°C for 20 minutes and blocked in 2% Boehringer blocking reagent, 20% heat-inactivated sheep serum in post-hyb for 1h. Probe was detected using alkaline phosphatase-labelled anti-digoxigenin antibody (1:1000, Promega, 11093274910) in blocking buffer at 4°C overnight. Sections were then washed with in post-hyb and equilibrated for 10 min in NTMT buffer (100 mM NaCl, 100 mM Tris HCl pH 9.5, 50 mM MgCl2, 1% Tween 20). Finally, sections were incubated with alkaline phosphatase substrates nitro blue tetrazolium (NBT) and 5-bromo-4-chloro-3-indolyl phosphate (BCIP) in NTMT buffer to reveal signal, washed in PBS-0.1% Triton X-100 and post-fixed in 4% PFA at 4°C and washed again in PBS-0.1% Triton X-100 and slides mounted with ProLong® Gold Antifade Mountant. Images were acquired on a Leica DMRB microscope fitted with a Nikon D7100 camera.

### TdT-UTP nick end labelling (TUNEL) assay

The inter-limb spinal cord tissue from CD1 mouse embryos was dissected and fixed at 4°C for 2 hours or overnight in 4% PFA. TUNEL assay was performed on 20 μm thick cryosections using ApopTag® Peroxidase In Situ Apoptosis Detection Kit (Merck Millipore, Cat. # S7100) according to manufacturer’s instructions.

### Tissue measurements and cell counts

Images were processed using Fiji and assembled using Adobe Photoshop and Illustrator software. Measurements of the dorsal-ventral length and apical-basal width of the ventricular layer were determined using Deltavision SoftWorx software. The widest point was taken as the measure for width. The Cell Counter plugin in Fiji was used to count SOX2 expressing cells labelled with pH3 or PCNA and the same approach was used to count fluorescent cells in tissues from the FUCCI mouse line.

### Statistical analysis

All the results were summarised as mean ± standard error of the mean (SEM). A two-sided, unpaired Student’s t-test with pooled/equal variance was used to test whether the differences in the means of two groups were statistically significant. A p-value below 0.05 was considered to be statistically significant. The number of samples studied in each experiment is mentioned in their respective figure legends and see Tables S3-S6. No statistical method was used to determine the sample size.

## Results

### Changes in tissue dimensions and local cell rearrangements are associated with ventricular layer attrition during formation of mouse spinal cord central canal

To characterise the changing dimensions and organisation of the ventricular layer in the developing mouse spinal cord, we delimited this cell population using immunofluorescence to detect SOX2-positive neural progenitors and surrounding TUJ1-expressing neurons. Measurements were made from embryonic day (E)12.5 to E18.5 in the thoracic spinal cord (Figs 1A-G’ and Table S3).

**Figure 1.**
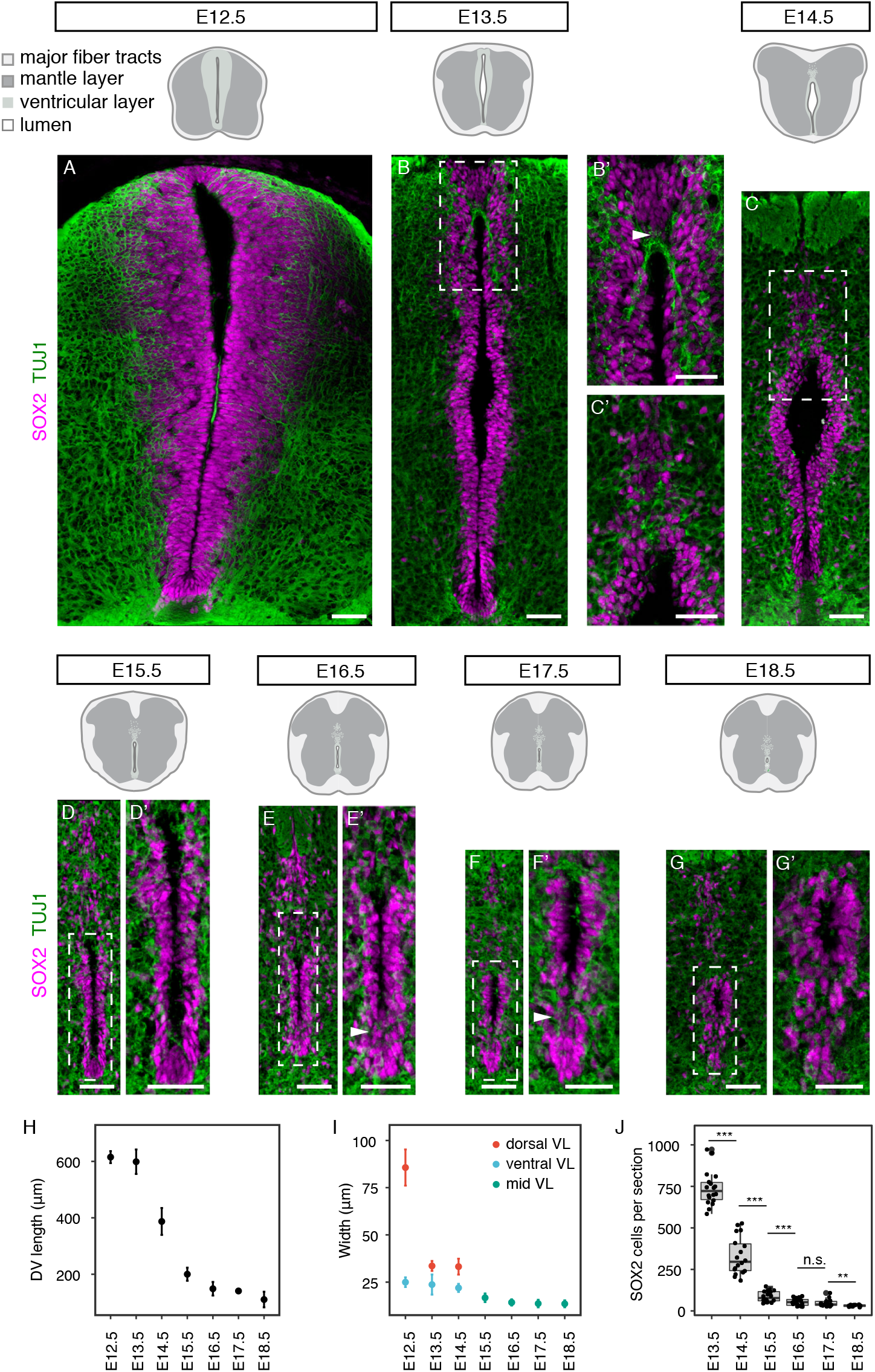
Changing ventricular layer dimensions during mouse spinal cord development. (A-G’) Immunofluorescence for SOX2 (magenta) and TUJ1 (green) reveals the changes in the organisation of the ventricular layer and the surrounding mantle layer (black dashed boxes) at the indicated stages. Higher magnification images of white dashed boxes from E13.5 to E18.5 are shown in B’, C’, D’, E’, F’ and G’, respectively. Note that some of the ventral-most SOX2-expressing cells appear to dissociate from the ventricular layer at later stages (arrowheads in E’ and F’). (H-I) Ventricular layer length (H) and width (I) at indicated stages (at least 9 sections, n = 3 embryos for each stage). Widest horizontal distance across the dorsal region of the ventricular layer (dorsal VL) and adjacent to the floor plate (ventral VL) was measured. From E15.5 the width of the ventricular layer becomes fairly uniform along the dorsal-ventral axis, so measurement was at the mid ventricular layer (mid VL) at these stages; (J) SOX2 positive cells per section were quantified at each stage (see Table S3). Error bars represent mean values ± standard error of the mean (SEM). A two-sided unpaired Student’s t-test was used to determine whether differences in the mean values were statistically significant, ***, *p* < 0.001; **, *p* < 0.01; * *p* < 0.05. Scale bars: 40 μm.

At E12.5, the length of the ventricular layer spans the height of the transverse section of the spinal cord (615.2 ± 7.1 μm) and it is widest dorsally, tapering ventrally (Figs 1A, H, I). At this stage, the apical surface of the neuroepithelium often meets at the midline, obscuring the central lumen in all but the dorsal region (Fig 1A). The length of the ventricular layer is unchanged at E13.5; however, its dorsal width is now substantially reduced (from 85.6 to 33.6 μm or from 10 to 4 cell diameters, Figs 1B, H, I). At this stage, the ventricular layer has an elongated rhomboid shape with the lumen open centrally (Fig 1B). Here, in a nexus about five cell diameters below the roof plate, TUJ1-positive neuronal processes are now visible at the dorsal midline and interdigitate between SOX2-positive cells, which have lost their contiguous pseudostratified arrangement (Figs 1B’). This local remodelling of the ventricular layer appears to be the first indication of the process of dorsal collapse.

By E14.5, the length of the ventricular layer is drastically reduced (to 387.1 ± 15.9 μm) due to dorsal collapse (Figs 1C, H). Almost all of the dorsal region now contains interdigitated neuronal processes and more sparsely distributed SOX2-positive cells (Fig 1C’). Some of these more dorsal SOX2 expressing cells have now come to abut the open central lumen, which has acquired an inverted tear-drop shape (Fig 1C’). This morphology, however, is quickly lost as dorsal collapse proceeds. Indeed, by E15.5 the ventricular layer is further reduced in length (200.1 ± 7.3 μm; or 15 to 20 cell diameters) and width, which becomes more uniform along the dorsal-ventral axis (16.7 ± 2.2 μm; or 2 cell diameters), and only a narrow slit-like lumen is apparent (Figs 1D, D’, H, I). At E16.5, the length and width of the ventricular layer continues to diminish (Figs 1E, E’, H, I) and some of the ventral-most SOX2-positive cells appear to dissociate from the ventricular layer (arrowheads in Figs 1E’). Between E17.5 and E18.5, the ventricular layer is reduced in length just a few more cell diameters, as the more loosely arranged SOX2-positive cells both dorsally and ventrally are further dispersed (110.4 ± 8.7 μm in length) (Figs 1F-G’, H). The remaining contiguous SOX2-expressing cell population that constitutes the ventricle appears now to have reached the final, oval-shaped arrangement of what will be the future spinal cord central canal (Figs 1G, G’).

These data identify large scale morphological changes and local cell re-arrangements associated with ventricular layer attrition. These include, first, a reduction in dorsal width at E13.5 accompanied by a local loss of pseudostratified epithelial organisation, followed by dramatic reduction in length associated with widespread loss of epithelial organisation in the dorsal region and so dorsal collapse at E14.5. This is reflected in a significant reduction in the number of SOX2-positive cells in the ventricular layer between E13.5 and E14.5 and between E14.5 and E15.5 (Fig 1J). In addition, a later further attrition step, which we name here ventral dissociation, separates most ventral SOX2-positive cells from the future central canal.

### Reduced cell proliferation, but not apoptosis, accompanies ventricular layer attrition

The reduction of SOX2-positive cells in the ventricular layer as development proceeds is at least in part accounted for by the delamination of cells from this proliferative zone to form neurons and, after E12.5, glial cells (Deneen et al., 2006, Stolt et al., 2003). In addition, it is also possible that progenitor cells remaining in the ventricular layer reduce their proliferation rate and so cell replacement slows, accelerating the reduction of this progenitor cell population. To test the latter possibility, we set out to investigate cell proliferation parameters in the ventricular layer from E13.5 to E18.5. As it is largely the ventral half of the ventricular layer at E13.5 that is retained following dorsal collapse, we focused on this cell population and on the whole contiguous SOX2-positive cell population from E14.5 onward (Figs 2Aa-F). We first used the punctate pattern of proliferating cell nuclear antigen (PCNA), which indicates sites of DNA synthesis (Bravo and Macdonald-Bravo, 1985), to determine the percentage of SOX2-positive cells in S phase at each stage. This analysis revealed a reduction in the proportion of cells in S phase between E13.5 to E14.5 (Figs 2Aa-C) and that thereafter this remains low and approximately constant to E18.5 (Fig 2C). Next, the mitotic index was calculated by determining the percentage of phospho-Histone H3 (pH3)-positive SOX2 cells at each stage. This revealed a similar pattern, in that the percentage of mitotic cells is low at E13.5 and shows a tendency to decrease further at later stages (the mean differences between each two consecutive stages is not statistically significant, but the mitotic index at E18.5 is significantly lower than that at E13.5, *p* = 0.017) (Figs 2Da-F).

**Figure 2.**
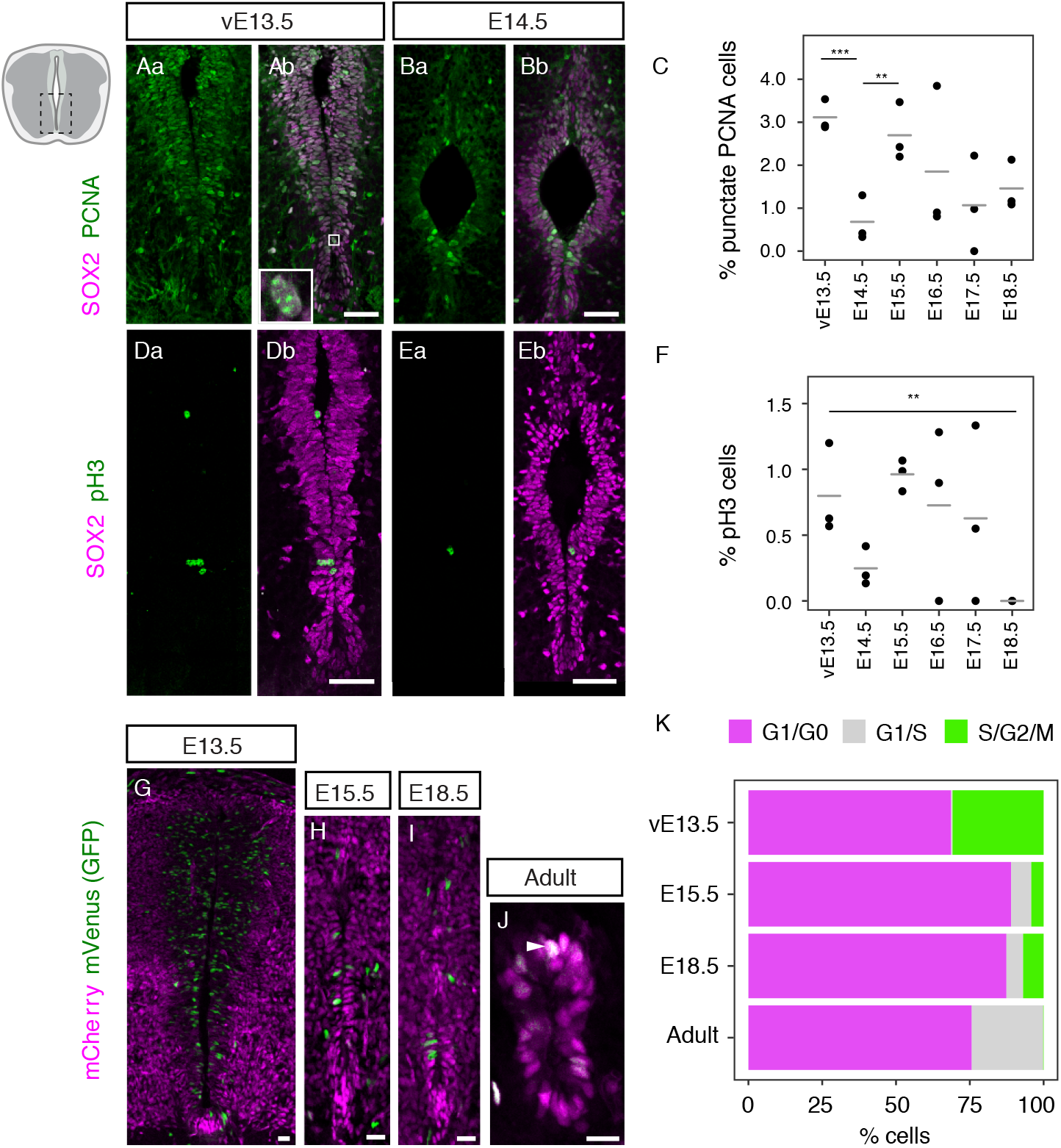
Cell proliferation declines as the spinal cord central canal forms. (Aa-Ab) Immunofluorescence for the neural progenitor marker SOX2 (magenta) and the cell proliferation marker PCNA (green) in the ventral half of the ventricular layer at E13.5 and (Ba-Bb) at E14.5. Only cells with a punctuate PCNA pattern were counted in the analysis (white full box in Ab); (C) Percentage of SOX2-postive cells with a punctate PCNA pattern in the spinal cord at E13.5 to E18.5. Each dot represents one embryo and the horizontal bars are the mean values of each stage; (Da-Db) Immunofluorescence for SOX2 (magenta) and the mitotic marker pH3 (green) in the ventral half of the ventricular layer at E13.5 and (Ea-Eb) E14.5; (F) Percentage of SOX2-postive cells labelled with mitotic marker pH3 in the spinal cord at E13.5 to E18.5. Each dot represents one embryo and the horizontal bars are the mean values of each stage; (G) mCherry-hCdt1 (magenta) and mVenus-hGeminin (green) in the ventricular layer at E13.5, (H) E15.5, (I) E18.5 and (J) the adult central canal; (K) Percentage of cells in G1/G0 (magenta), G1/S (grey), and S/G2/M (green) in the ventral half of the ventricular layer at E13.5, the whole ventricular layer at E15.5 and E18.5, and in the adult central canal of Fucci2a mice. Asterisks show significant results from two-sided, unpaired Student’s t-tests between each two correlative stages. ***, *p* < 0.001; **, *p* < 0.01; \**p* < 0.05. Only statistically significant differences are indicated. The number of cells and sections analysed for each embryo and the exact p-values for all comparisons can be found in Table S4 (for C), Table S5 (for F), and Table S6 (for K). Scale bars Aa-Eb: 40 μm, G-J: 20 µm.

A limitation of the above approaches is the small number of cells observed with a punctate PCNA pattern or undergoing mitosis (Table S3). To extend this cell proliferation analysis, we therefore next assessed cell cycle phase distribution in ventricular layer cells using Fucci2a transgenic mice (Mort et al., 2014) at key stages: before (ventral E13.5) and after (E15.5) dorsal collapse, after ventral dissociation (E18.5), and in the adult central canal (Figs 2G-J). These transgenic mice express two cell cycle-regulated proteins fused to fluorescent proteins, one to mCherry (mCherry-hCdt1) and the other to mVenus (mVenus-hGeminin). This results in the accumulation of mCherry-hCdt1 during the G1 and G0 phases of the cell cycle, making cells in G1 and G0 fluoresce red (shown here in magenta). mCherry-hCdt1 is degraded at the G1/S transition following interaction with Geminin (Wohlschlegel et al., 2000, Tada et al., 2001) and mVenus-hGeminin begins to accumulate in S, so cells in the G1/S transition fluoresce both red and green and appear yellow (shown here in white). mVenus-hGeminin continues to be expressed during S/G2/M phases, making cells in these phases fluoresce bright green. mVenus-hGeminin is then rapidly degraded before cytokinesis (Mort et al., 2014, Sakaue-Sawano et al., 2008). In agreement with our previous measurements, we found that the percentage of cells in S/G2/M decreases after E13.5 (from 30.8 ± 4.7% to 4.2 ± 0.5% E15.5, mean ± SEM, *p* = 0.027) and remains relatively constant to E18.5 (6.9 ±1.6%, *p* = 0.239) (Fig 2K). In the adult central canal, cells in S/G2/M are rarely detected (0.1 ± 0.1%, *p* = 0.003). In contrast, the percentage of cells in the G1/G0 state increase from E13.5 to E15.5, from 68.7 ± 4.8% to 89.0 ± 1.9% (*p* = 0.06), although the difference is not statistically significant. The percentage of cells in G1/G0 remains rather similar between E15.5 and E18.5 (87.4 ± 2.2%, *p* = 0.39). This may reflect the intriguing emergence a group of cells transitioning from G1 to S, which becomes first evident at late developmental stages (from 0.48 ± 0.24% at E13.5 to 6.8 ± 2.0% at E15.5, *p* = 0.006) and increases still further in adult central canal cells (from 5.6 ± 1.6% at E18.5 to 24.4 ± 1.3%, *p* = 0.0004) (Fig 2K). This suggests that while the majority of adult central canal cells reside in G1/G0, the rest of this cell population now appear poised to progress through the cell cycle as they are beginning to upregulate Geminin (early S-phase) and lose Cdt1 (Figs 2J, K) (see Discussion). These findings show that after E13.5 gradually fewer cells progress through S/G2/M phases of the cell cycle and instead spend longer in G1. Our analysis suggests that this involves either arrest as cells exit the cell cycle into G0 or G1 lengthening, with about a quarter of adult cells being in a G1/S phase transition state. Together, these changes indicate a reduction in cell proliferation in the spinal cord ventricular layer as it forms the central canal.

A further cellular process that could contribute to the attrition of the ventricular layer is localised programmed cell death (apoptosis). This might operate in the dorsal region as the collapse commences from E13.5 and/or during subsequent refinement of the ventricular layer cell population as the central canal forms. To investigate this possibility, we carried out a TdT-UTP nick end labelling (TUNEL) assay on transverse sections between E12.5 and E18.5 to detect DNA fragmentation, a hallmark of apoptosis. This revealed few apoptotic cells at any stage (Fig S1A-G) and this was confirmed by immunofluorescence for activated caspase-3 (Fig S1H-O). These findings indicate that apoptosis does not contribute to ventricular layer attrition.

### Dissociation of a ventral-most cell population contributes to central canal formation

Analysis of the ventral-most SOX2-positive cells during development revealed a spatial rearrangement from E15.5, which culminated in the dissociation of the majority of these cells from the forming central canal (Figs 1D-G’). To define further this ventral cell rearrangement and the cell populations involved, we analysed expression of marker proteins of the ventral-most cell population, the floor plate, as the central canal forms. FOXA2 is a transcription factor that is initially localised in the floor plate and the adjacent p3 interneuron domain (Fig 3A) (Cruz et al., 2010, Schafer et al., 2007). We found that this pattern of expression persists until E14.5 (Fig 3B) and from E15.5, while ventral ventricular layer cells remain FOXA2-positive, the ventral-most cells begin to dissociate into a distinct group (arrowheads in Figs 3C-F). In addition, a new group of cells that express FOXA2 highly, now also appears sub-ependymally (outside the ventricle) (Figs 3C-F). Moreover, while FOXA2 expression later is no longer detected in the adult central, these FOXA2-high sub-ependymal cells remain (Fig 3G) and may correspond to a subset of cerebrospinal fluid-contacting neurons (CSF-cN”) (Petracca et al., 2016). *Arx*, is a further well-known floor plate marker from E9 to E12.5 (Miura et al., 1997, Cruz et al., 2010). A group of ARX-positive cells continued to define this cell population at E13.5 (Fig 3H). At later stages, ARX-positive cells form an elongated arrangement, again with some cells remaining in the ventral ventricular layer while others dissociate to form a distinct ventral group (arrowheads in Figs 3I-L). In contrast with FOXA2 expression, some ARX-positive cells then persist in the adult ventral central canal, as noted previously (Khazanov et al., 2017) (Fig 3M). These patterns of expression corroborate the dissociation of ventral-most floor plate cells observed by monitoring changing arrangement of SOX2-expressing cells and support the inclusion of some floor plate cells in the adult central canal.

**Figure 3.**
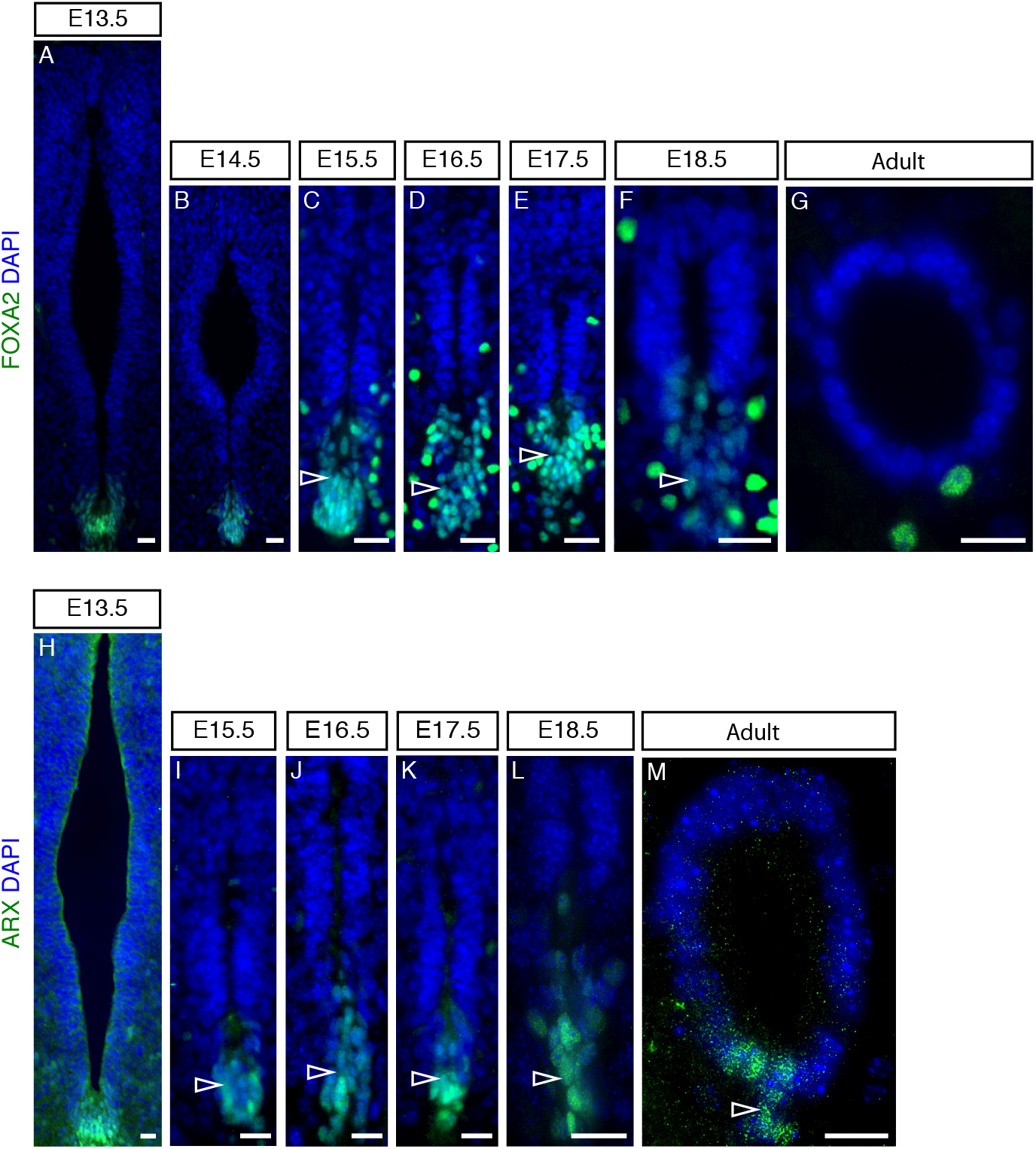
FOXA2 and ARX expression in the developing and adult mouse spinal cord. (A-G) Immunofluorescence for FOXA2 (green) at the indicated stages (E13.5: 33 sections, n = 3 embryos; E14.5: 8 sections, n = 2 embryos; E15.5: 18 sections, n = 5 embryos; E16.5: 22 sections, n = 5 embryos; E17.5: 16 sections, n = 4 embryos; E18.5: 26 sections, n = 4 embryos; adult (10 – 11 week old mice): 18 sections, n = 6 mice); (H-M) Immunofluorescence for ARX expression (green) at the indicated stages (E13.5: 12 sections, n = 1 embryo; E15.5: 10 sections, n = 2 embryos; E16.5: 10 sections, n = 1 embryo; E17.5: 11 sections, n = 1 embryo; E18.5: 12 sections, n = 2 embryos; adult (10 and 24 week old mice): 5 sections, n = 2 mice each stage. Dorsal is top, ventral is bottom. Nuclei are stained with DAPI (blue). All scale bars: 20 μm.

To elucidate further the remodelling of this ventral cell population, we next combined FOXA2 and TUJ1 immunofluorescence, to visualise the relationship between FOXA2-positive cell groups and neuronal processes. This revealed TUJ1-positive cell processes intervening between the central canal and the dissociating ventral-most cell group from E15.5, with a pronounced ventral commissure apparent by E18.5 (Figs 4Aa-Dd).

**Figure 4.**
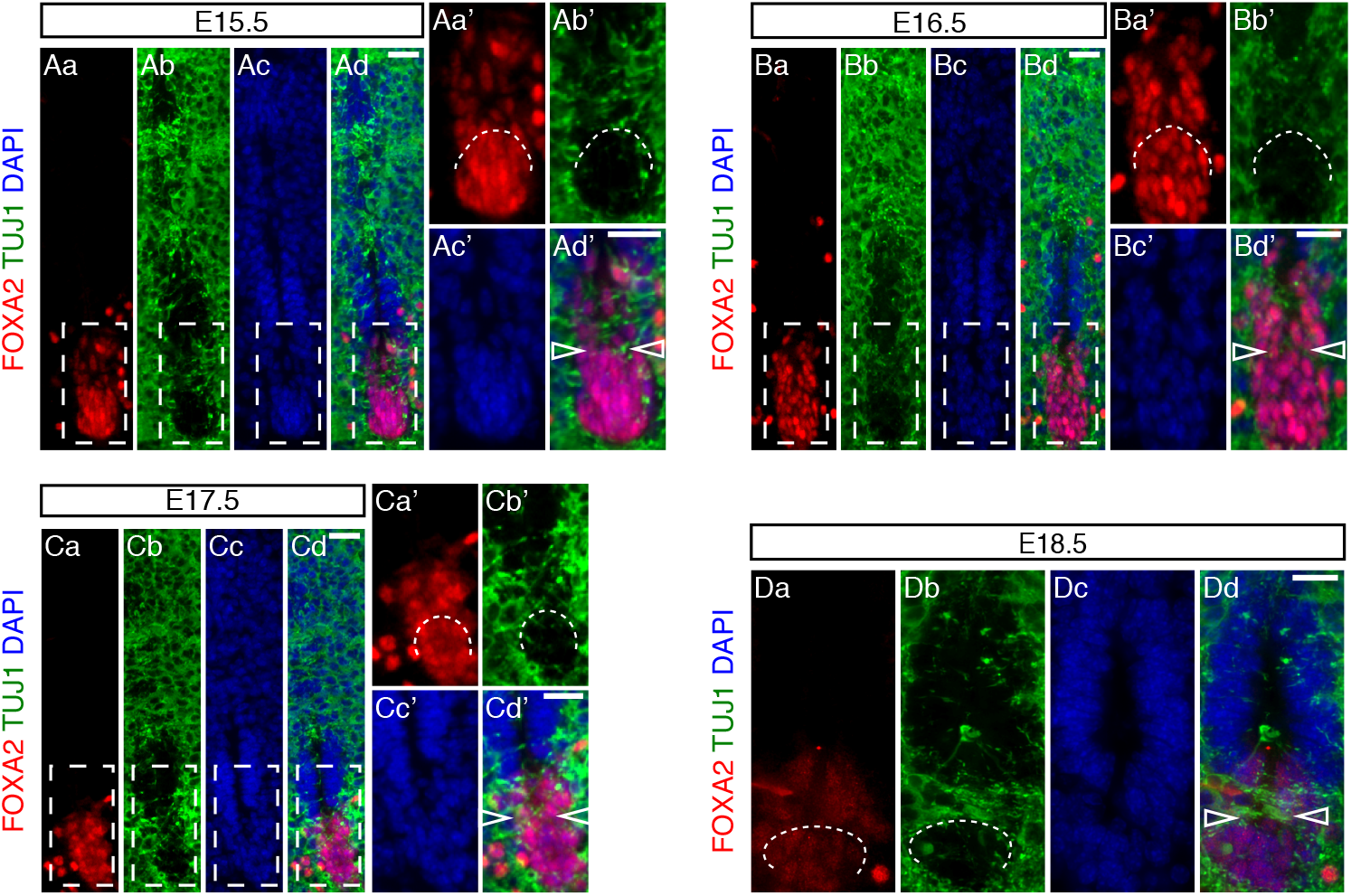
Ventral cell rearrangement during late mouse spinal cord development. (Aa-Dd) Immunofluorescence for FOXA2 (red) and TUJ1 (green) in mouse spinal cord development at stages indicated (E15.5: 11 sections, n = 3 embryos; E16.5: 10 sections, n = 3 embryos; E17.5: 9 sections, n = 3 embryos; E18.5: 9 sections, n = 2 embryos). Higher magnification of regions outlined in white dashed line boxes are shown in Aa’-Ad’, Ba’-Bd’ and Ca’-Cd’. Partition of the ventral-most floor plate cell population by TUJ1-positive neuronal processes, indicated by curved white dashed lines and by arrowheads in merged images. Nuclei are stained with DAPI (blue). All scale bars: 20 μm.

Intriguingly, this dissociation of the ventral-most cell population occurs concomitant with a change in the expression pattern of *Foxj1*, the master regulator of the ciliogenesis programme (Choksi et al., 2014) and a mediator of migratory cell behaviour of spinal cord ependymal cells in response to injury (Li et al., 2018). *Foxj1* transcript and protein levels decline in the floor plate as ventral dissociation begins at E15.5 (compare ventral region in Figs 5A-D and 5Ha-Ib), but is upregulated in ventral progenitors from E14.5 (Figs 5C-5G and 5Ha-5Lb) and expressed in the majority of central canal cells, although only at low-level in ventral-most cells at E18.5 (Li et al., 2018). Together, these observations identify a distinct dissociation event which resolves the ventral ventricular cell population into two anatomically distinct groups, the ventral central canal and the dissociated ventral-most floor plate.

**Figure 5.**
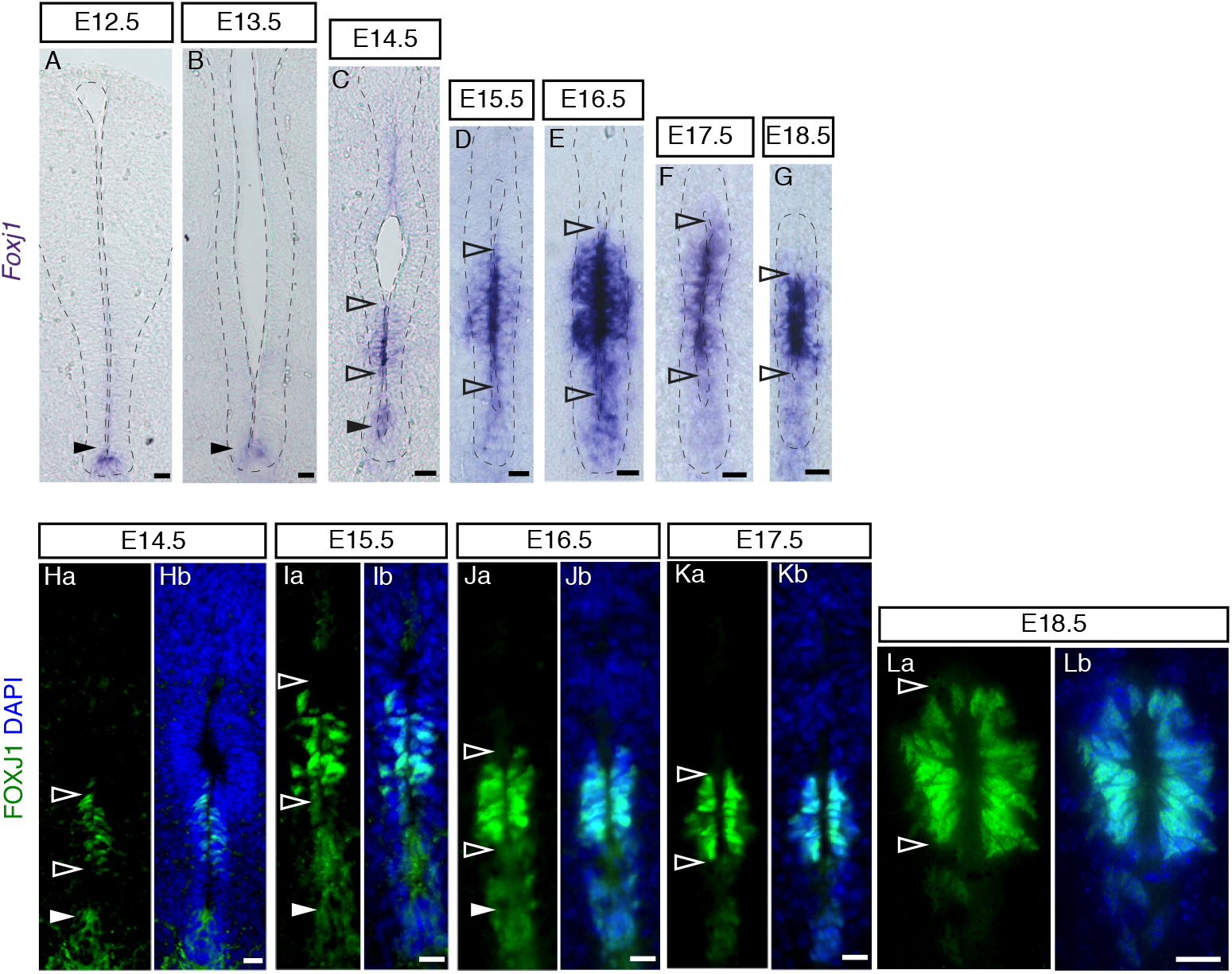
Foxj1 mRNA and protein expression during mouse spinal cord development. (A-G) *In situ* hybridisation showing the dynamic expression of *Foxj1* mRNA at the indicated stages (E12.5: 22 sections, n = 5 embryos; E13.5: 32 sections, n = 7 embryos; E14.5: 32 sections, n = 7 embryos; E15.5: 33 sections, n = 12 embryos; E16.5: 86 sections, n = 15 embryos; E17.5: 114 sections, n = 18 embryos; E18.5: 94 sections, n = 11 embryos). Note apical localisation of transcripts. Nuclei are stained with DAPI (blue); (Ha-Lb) Immunofluorescence showing dynamic expression of the FOXJ1 protein (green) at indicated stages (E14.5: 66 sections, n = 4 embryos; E15.5: 33 sections, n = 3 embryos; we also confirm findings of Li et al 2018 at later stages E16.5: 12 sections, n = 1 embryo; E17.5: 12 sections, n = 1 embryo; E18.5: 12 sections, n = 1 embryo). Nuclei are stained with DAPI (blue). Note that low levels/gaps in expression are found in the ventricular layer between the region closer to the dorsal lumen and the domain adjacent to the floor plate at E14.5 (arrowheads in C and Ha). Scale bars for A, B: 40 μm, for C-Lb: 20 μm.

### A floor plate contribution to the spinal cord central canal is supported by analysis of Shh pathway components, which also reveals a gradual decline in Shh signalling

The presence of ARX-positive cells in the ventral central canal supports the notion that some floor plate-derived cells contribute to this structure. Indeed, lineage tracing of cells expressing the floor plate marker *Nato3* (*Ferd3l*), using a Nato3-LacZ line, revealed some LacZ-positive cells in the spinal cord central canal (Khazanov et al., 2017). Further, the regulatory relationships between *Foxa2*, *Nato3* and Shh signalling that underpin floor plate maturation (Cruz et al., 2010, Mansour et al., 2014, Mansour et al., 2011), suggest that the Shh pathway is active as the central canal forms (Rowitch et al., 1999). Moreover, as Shh acts as a mitogen as well as a morphogen in the early neural tube it is interesting to assess whether cells retaining floor plate markers continue to act as a signalling centre during mid-late development and postnatally.

Using *in situ* hybridization, we found that transcripts for *Shh* in the floor plate were attenuated at E15.5 and undetectable by E16.5 (Figs 6A-D). In contrast, transcripts for the Shh receptor and transcriptional target *Ptch1* (Goodrich et al., 1997, Marigo and Tabin, 1996, Pearse et al., 2001, Vokes et al., 2008) persisted for longer in the presumptive central canal, but were much reduced by E18.5 (Figs 6E-J). These observations suggest that Shh signalling persists after *Shh* transcription ceases, but declines as the central canal forms. We investigated this further by looking at GFP expression in two Shh signalling reporter mouse lines: one expressing a tagged Shh-GFP transgene (Chamberlain et al., 2008) and another expressing Gli-binding site motifs linked to GFP (GBS-GFP), which provides a further readout of Shh signalling activity (Balaskas et al., 2012). Expression of Shh-GFP was restricted to the floor plate at all embryonic stages examined including E18.5 (Figs 6Ka-Mb). This persistence of Shh-GFP compared with that of *Shh* transcripts may be indicative of residual Shh protein (consistent with continued *Ptch1* expression), but could also reflect a longer half-life of the GFP tagged protein. Ependymal cells of the adult central canal, however, lacked any Shh-GFP (Fig 6N).

**Figure 6.**
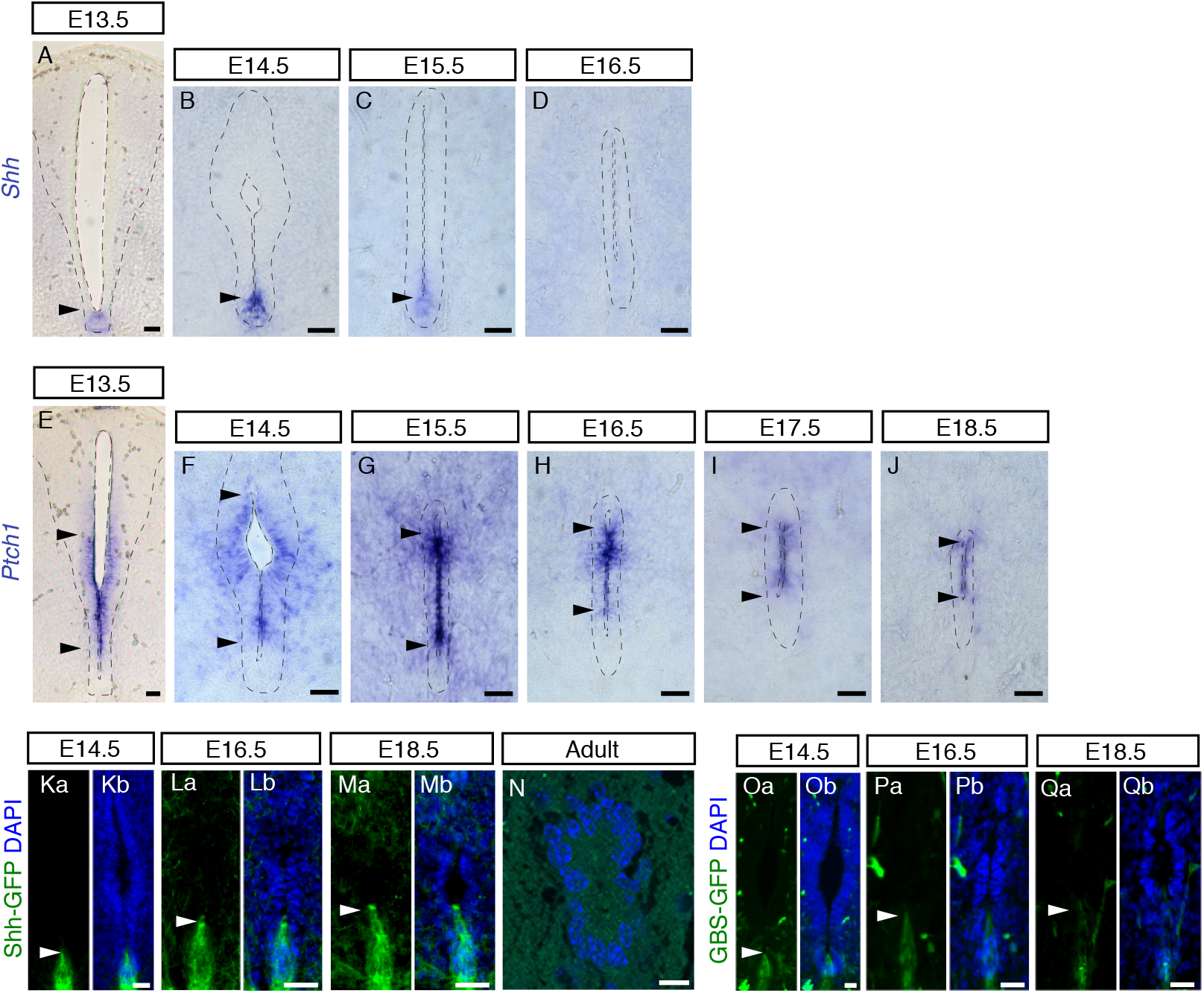
Shh activity persists after decline in *Shh* transcripts during mouse spinal cord development. (A-D) *In situ* hybridisation showing *Shh* mRNA expression in the floor plate (arrowheads) at the indicated stages (E13.5: 14 sections, n = 3 embryos; E14.5: 15 sections, n = 5 embryos; E15.5: 58 sections/explants, n = 9 embryos; E16.5: 38 sections/explants, n = 9 embryos). From E17.5 onwards, *Shh* mRNA expression was no longer detected in the spinal cord (E17.5: 38 sections/explants, n = 8 embryos; E18.5: 18 sections, n = 3 embryos; 24- and 40-week-old: 17 sections, n = 3 adult mice; data not shown); (E-J) *Ptch1* mRNA expression at the indicated stages (E13.5: 27 sections, n = 5 embryos; E14.5: 26 sections/explants, n = 5 embryos; E15.5: 43 sections/explants, n = 6 embryos; E16.5: 49 sections/explants, n = 8 embryos; E17.5: 40 sections/explants, n = 8 embryos; E18.5: 16 sections, n = 3 embryos). Note *Ptch1* is excluded from the floor plate and detected in ventral half of the ventricular layer (between arrowheads) and persists in the forming central canal E18.5, but was no longer detected in adult (40-week-old: 24 sections, n = 3 mice; data not shown). (Ka-Mb) Immunofluorescence for GFP in Shh-GFP embryos was detected in the floor plate during development (arrowheads) E14.5: 17 sections, n = 1 embryo; E16.5: 17 sections, n = 1 embryo; E18.5: 18 sections, n = 1 embryo, (N) but not in adult central canal (9-10 week adult (88 sections, n = 3 mice); (Oa-Qb) Immunofluorescence for GFP (arrowheads) in GBS-GFP embryos revealed positive cells in the ventral region at indicated stages (E14.5: 47 sections, n = 3 embryos; E16.5: 43 sections, n = 3 embryos; E18.5: 33 sections, n = 3 embryo). Ka-Qb, nuclei are stained with DAPI (blue). All scale bars: 40 μm, except N: 10 μm.

The presence of cytoplasmic GFP in these Shh-GFP embryos further allowed us to observe changing cell morphologies and confirmed the elongation of cells associated with the ventral dissociation event (Figs 6Ka-Lb). Moreover, some GFP-expressing cell processes remain integrated within the apical surface of the ventricle even at late stages (Fig 6Ma-Mb); this may reflect inclusion of floor plate-derived cells in the central canal or additional retained contact of the ventral-most dissociated cell group.

GBS-GFP was expressed more widely than Shh-GFP and was detected in the floor plate and adjacent ventricular layer cells at E14.5 (Fig 6Oa-Ob). However, it too also became restricted to the floor plate at subsequent stages, with a dwindling number of expressing cells continuing to contact the ventricular lumen at E18.5 (Figs 6Oa-Qb). These data indicate that Shh signalling persists locally for some days after *Shh* transcription ceases at E14.5, declining slowly as the central canal forms and appears lost in the adult. The continued contact of floor plate-derived cells with the apical surface of central canal lumen, however, further substantiates the contribution of some cells of floor plate origin to this structure.

### Dorsal BMP signalling persists and extends ventrally during central canal formation

The roof plate origin of dorsally located radial glia-like cells that remain attached to the central canal during dorsal collapse was recently confirmed in mice (Xing et al., 2018, Shinozuka et al., 2019). During development, roof plate cells express Wnt and BMP ligands, which act in opposition to Shh signalling (Le Dreau and Marti, 2012). Moreover, BMP signalling has been implicated in the regulation of quiescence and proliferation of adult brain neural stem cells (Llorens-Bobadilla et al., 2015, Mira et al., 2010).

To investigate how BMP signalling changes as the central canal forms, we carried out immunofluorescence against phosphorylated SMAD1/5 (pSMAD1/5), a readout of BMP signalling, and the BMP receptor BMPR1B. Analysis of patterns of pSMAD1/5 detection revealed that BMP activity is restricted to roof plate cells at E12.5 and persists in SOX2-positive cells located dorsally until at least E16.5 (Figs 7Aa-Db). However, from E13.5 pSMAD1/5 was additionally detected in a ventral domain (Fig 7Ba-Bb) which became more widespread at E14.5 (Fig 7Ca-Cb) and continued to be detected dorsally and ventrally at E16.5 (Fig 7Da-Db). Analysis of BMPR1B expression revealed that this was present in the roof plate at E12.5 (Fig 7Ea-Eb) and that this persisted and extended ventrally at E16.5, where the protein was enriched apically (Fig 7Fa-Fb). This pattern of expression continued in the adult central canal, with low level protein detected ventrally (Fig 7Ga-Gb). These findings indicate that BMP signalling is dynamically regulated in the developing spinal cord: initial dorsally-restricted activity spreads to ventral regions and this is mirrored in the changing expression pattern of BMPR1B.

**Figure 7.**
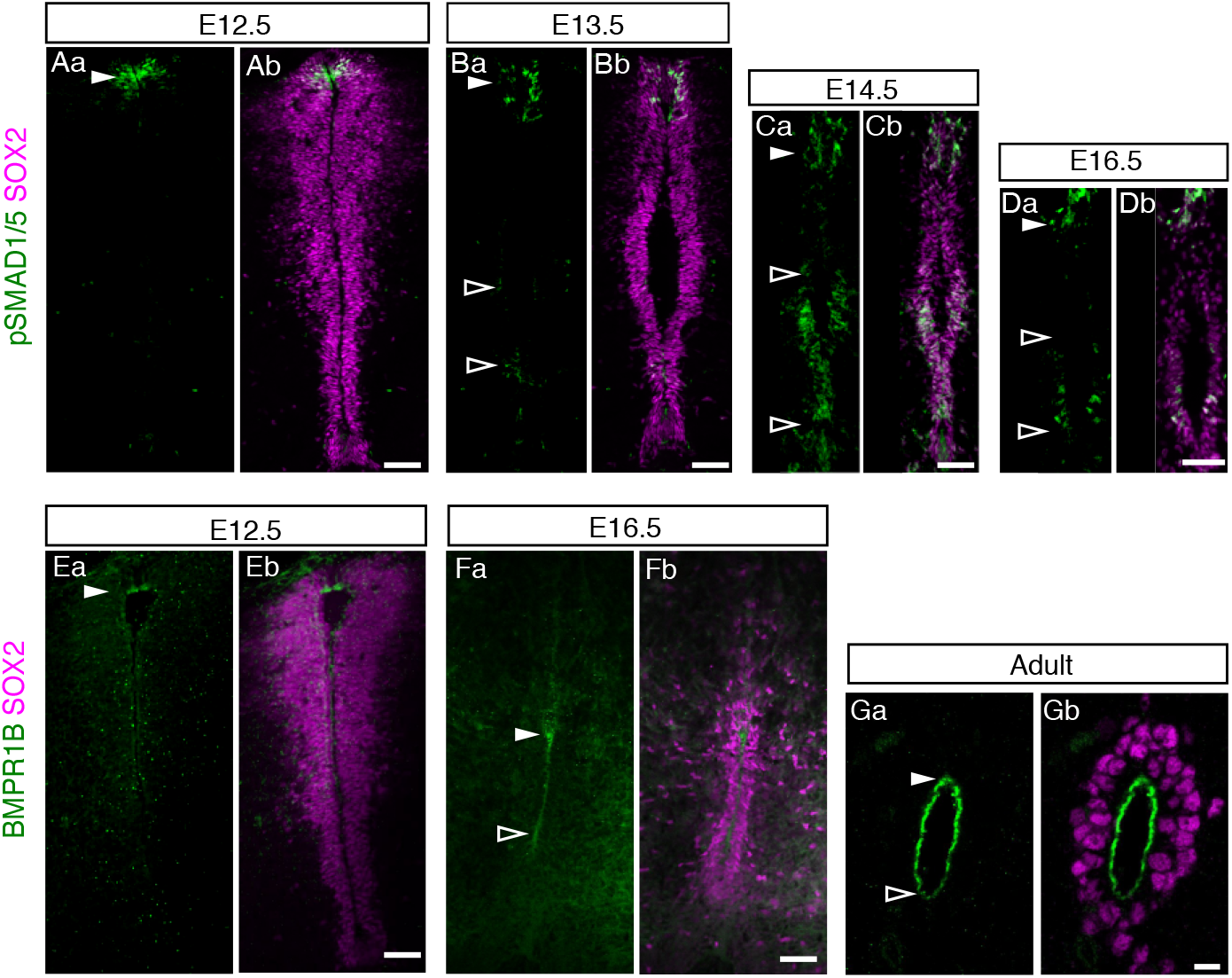
Expanding BMP signalling and BMPR1B expression during mouse spinal cord development. (Aa-Bb) Immunofluorescence for phospho-SMAD1/5 (green) detected in a dorsal cell population at E12.5 (16 sections, n = 2 embryos) and E13.5 (16 sections, n = 3 embryos) (full arrowheads), phospho-SMAD1/5 was also detected in the ventral ventricular layer at E13.5 (outlined arrowhead); (Ca-Db) At E14.5 (36 sections, n = 3 embryos) and E16.5 (14 sections, n = 2 embryos), phospho-SMAD1/5 was detected in dorsal (full arrowheads) and ventral regions of the ventricular layer (outlined arrowheads). (Ea-Fb) BMPR1B (green) was enriched dorsally at E12.5 (10 sections, n = 1 embryo) (full arrowhead) and was detected dorsally and apically in ventricle abutting cells (between arrowheads) at E16.5 (23 sections, n = 3 embryos); (Ga, Gb) Pronounced apical localisation of BMPR1B in adult central canal (10 – 11 week old mice: 22 sections, n = 2 mice), which was more weakly detected ventrally (outline arrowhead). Progenitor cells within the ventricular layer are labelled with SOX2 (magenta). All scale bars: 50 μm, except Adult: 10 μm.

## Discussion

This study provides evidence for multiple steps that contribute to the reduction of the spinal cord ventricular layer during development and lead ultimately to the formation of the adult spinal cord central canal. These include initial local loss of ventricular layer organisation below the roof plate, which spreads as dorsal collapse proceeds; a reduction in cell proliferation, involving an increase in the percentage of cells in the G1/S phase of the cell cycle; and cell rearrangement in the floor plate, involving dissociation of the ventral-most cell population. Despite this ventral remodelling, we provide further evidence that some floor plate cells are incorporated into the forming central canal. Overall, these data support a heterogeneous origin of the ependymal cell population that constitutes the adult spinal cord central canal, including roof plate and floor plate cells as well as ventral progenitor cells derived from *Pax6* and *Nkx6-1* expression domains.

We show that ventricular layer attrition begins not long after the switch from neurogenesis to gliogenesis in the developing spinal cord, and is first manifest by the narrowing and local disorganisation of the dorsal ventricular layer at E13.5. The disruption of the close alignment of SOX2-expressing progenitor cells below the roof plate appears to be the initiation site for dorsal collapse, which progresses rapidly in subsequent days. The appearance of neuronal cell processes between these progenitor cells and across the midline, may equate to the initiation of the dorsal commissure, which is more apparent at later stages in this position (Comer et al., 2015). It is interesting that the novel cell rearrangement we observe later in ventral-most regions from E16.5, named here as ventral dissociation, is also associated with the appearance of neuronal cell processes that traverse the midline, here to form the ventral commissure (Comer et al., 2015). In this case, these axons spatially separate most floor plate cells from cells that will form the adult central canal. This coincidence raises the possibility that commissure establishment is one of the mechanisms by which the ventricular layer is restricted in ventral as a well as dorsal regions (Fig 8).

**Figure 8.**
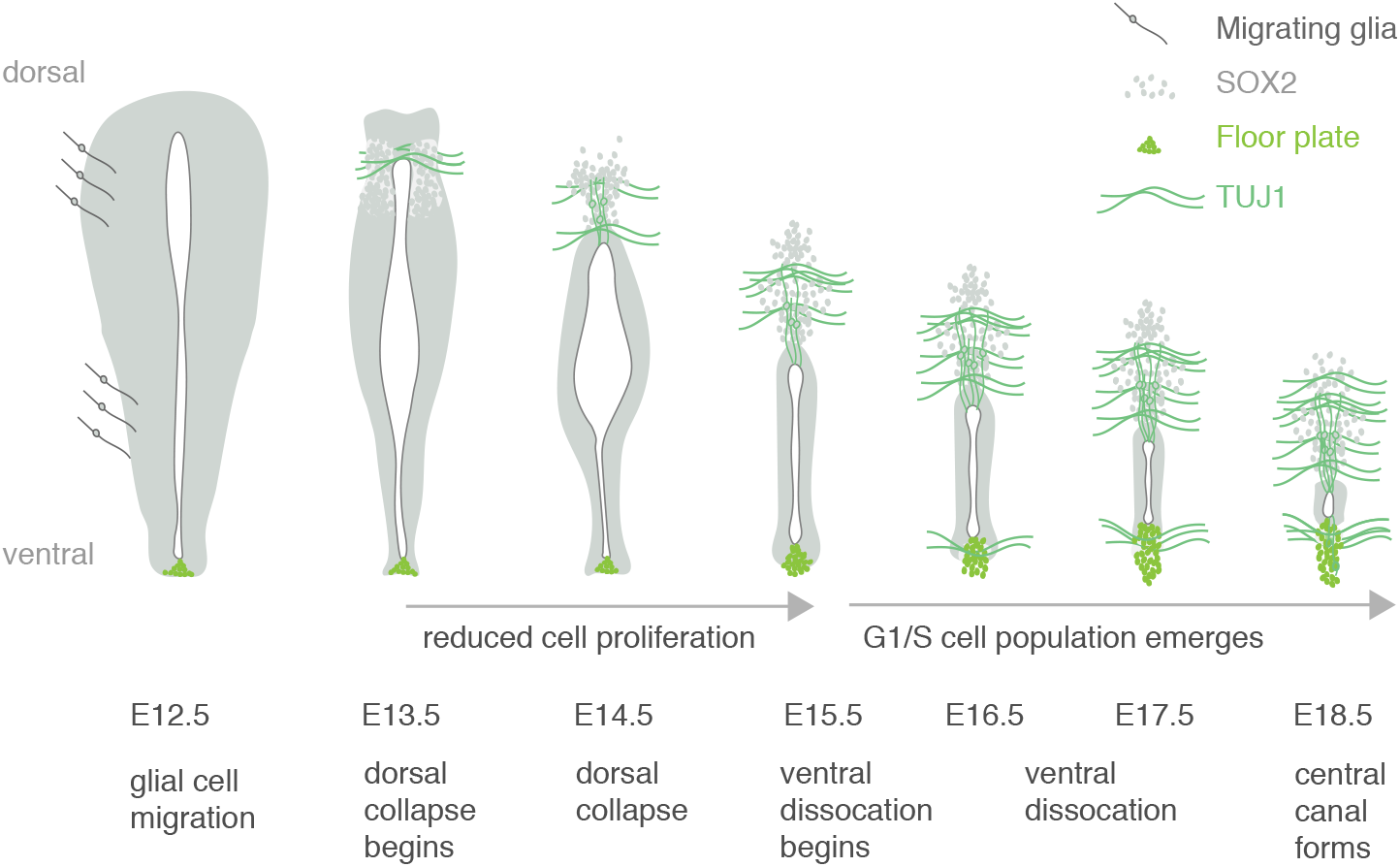
Summary of steps contributing to spinal cord ventricular layer attrition. Schematic of key changes in cell arrangement and cell cycle in the spinal cord ventricular layer at daily intervals leading up to the formation of central canal. Glial cell migration precedes dorsal collapse, which is accompanied by appearance of the dorsal commissure and reduced proliferation of ventricular layer cells. This is then followed by ventral dissociation of the floor plate cell population, accompanied by appearance of the ventral commissure and an increase in percentage of cells in G1/S phase of the cell cycle.

Studies at earlier stages in the mouse embryo (between E9.5 and E11.5) have shown that the cell proliferation rate in the ventricular layer declines during neurogenesis, with little difference between dorso-ventrally distinct progenitor cell populations (Kicheva et al., 2014). Our finding of a reduction in the proportion of cells in S and M phases of the cell cycle at later stages suggests that this decline in proliferation continues and contributes to ventricular layer attrition after E13.5. Intriguingly, our analysis of the cell cycle using Fucci2a reporter mice uncovered a progressive increase in cells in G1/S from E15.5, which ultimately accounted for almost a quarter of the cells in the adult central canal. As cells are asynchronously distributed across the cell cycle, this could be attributed to G1 lengthening: with more cells in G1 resulting in an increase in cells transitioning from G1 to S phase at a given time. Another intriguing possibility is that ventricular cells progressively switch to a cell cycle variant known as endocycle as they differentiate into ependymal cells. Endocycling cells go through G1 and S but do not undergo mitosis, becoming polyploid (Edgar et al., 2014). This would be consistent with the progressive increase in the percentage of G1/S cells and the decrease in S/G2/M cells we detect in Fucci2a embryos and mice. Indeed, in other studies using Fucci reporters, endoreplicating cells seem to co-express Geminin and Cdt1 (and thus appear yellow) for longer (Cao et al., 2017, Lazzeri et al., 2018). A switch to endocycling would potentially lead to higher genomic output, which might then enhance the secretory function of these cells. A further possibility is that the overall decline in cell proliferation is due to an increase in the cells that enter the quiescent state (G0) in G1. However, it is difficult to distinguish cells in G1 or G0 using the Fucci2a system as they both appear red. Whether the movement of cells into a reversible G0 state explains why adult central canal cells are mostly quiescent but can readily re-enter the cell cycle in response to multiple extrinsic stimuli or if G1/S cells also contribute to the reactivity of this cell population remains to be investigated. The latter are certainly an intriguing cell population that is not apparent in post-mitotic brain ependymal cells analysed in the Fucci2a mouse (Ford et al., 2018).

Early expression pattern studies in a range of amniote embryos suggest that the progenitor cell population expressing *Pax6* and *Nkx6-1*, but not *Nkx2-2*, is retained during development and gives rise to the adult central canal (Fu et al., 2003, Yu et al., 2013). Accumulating evidence, however, shows that the origin of spinal cord central canal cells is rather heterogeneous, including cells derived from the roof plate (Xing et al., 2018, Shinozuka et al., 2019) and from the floor plate (Khazanov et al., 2017, Ghazale et al., 2019) (and this study). Here, we found that cells expressing the floor plate marker ARX, but not FOXA2, are present in the adult spinal cord central canal and that Shh-GFP expressing cells are found integrated in the forming central canal at late embryonic stages.

An outstanding question is whether cells derived from the roof plate and floor plate remain as signalling centres during late embryonic development and into the adult spinal cord, as they do in regenerative species such as the axolotl and zebrafish (Schnapp 2005, Reimer 2009). Here, we have shown that Shh signalling declines from the ventral midline/floor plate at late embryonic stages and that Shh-GFP is no longer detected in central canal cells of the adult spinal cord. While it is possible that the detection of GFP in Shh-GFP and GBS-GFP transgenic embryos reflects the relatively long half-life of GFP rather than persistence of SHH, the detection of Shh targets *Ptch1* (this study) and *Gli* (Yu et al., 2013), support the possibility of dwindling Shh signalling at these late stages. This would also be consistent with studies using transgenic mice to delete *Shh* from the floor plate or its mediator *Smoothened* from the neural tube (Yu et al., 2013), as well as *Gli2* mutants (Ding et al., 1998, Matise et al., 1998), all of which fail to form a proper central canal, although confirmation of a persisting role for Shh signalling requires interference with this pathway specifically at late stages or in the adult.

In contrast, we found that the activity of the BMP signalling pathway, which opposes Shh signalling in the developing spinal cord, although initially dorsal expands as the central canal forms, with pSMAD1/5 detected extensively in more ventral regions as well as in dorsal roof plate-derived cells. This is further reflected by BMPR1B expression, which was detected in all cells in the adult central canal albeit at a lower level in the most ventral cells. A recent study has shown that Wnt signalling first from roof plate cells and later from the ependymal cells is required to maintain the adult central canal (Xing et al., 2018), while Wnt promotion of BMP signalling (found at earlier stages (Ille et al., 2007)), would be further consistent with the expansion of dorsal signalling. Taken together, this investigation of these key signalling pathways in the late embryonic and adult spinal cord suggests that dorsal signalling comes to prevail in cells of the ventricular layer as the central canal forms and ependymal cells differentiate. The involvement of BMP signalling in the regulation of adult neural stem cell quiescence in the brain (Llorens-Bobadilla et al., 2015, Mira et al., 2010) now prompts further experiments to investigate whether this spread of BMP activity promotes acquisition of the quiescent cell state, including changes in cell cycle and thereby sets aside the elusive spinal cord neural stem cell as the central canal forms.

## Acknowledgements

We thank Prof Ian Jackson and Richard Mort (IGMM, Edinburgh) for embryos and spinal cords from the Fucci2a mouse line, Dr James Briscoe (Francis Crick Institute) for mouse *Foxj1* cDNA and embryos and spinal cords from Shh-GFP and GBS-GFP transgenic mice, Prof Marysia Placzek (University of Sheffield) for mouse *Shh* cDNA, and Prof Benjamin Allen (University of Michigan, MI, USA) for the mouse *Ptch1* cDNA. We thank the Dundee Imaging facility for training and advice and members of the Storey group for comments on this manuscript.

## Author contributions

KGS and PF designed the project. MAC, GS, NS and MG designed and performed experiments. MAC, ARA and KGS interpreted the data and wrote the paper. All authors read and approved the final manuscript.

