## Supplementary material for "Multiple steps mediate ventricular layer attrition to form the adult mouse spinal cord central canal": Figure S1 & Tables S1 & S2

### Supplementary figures

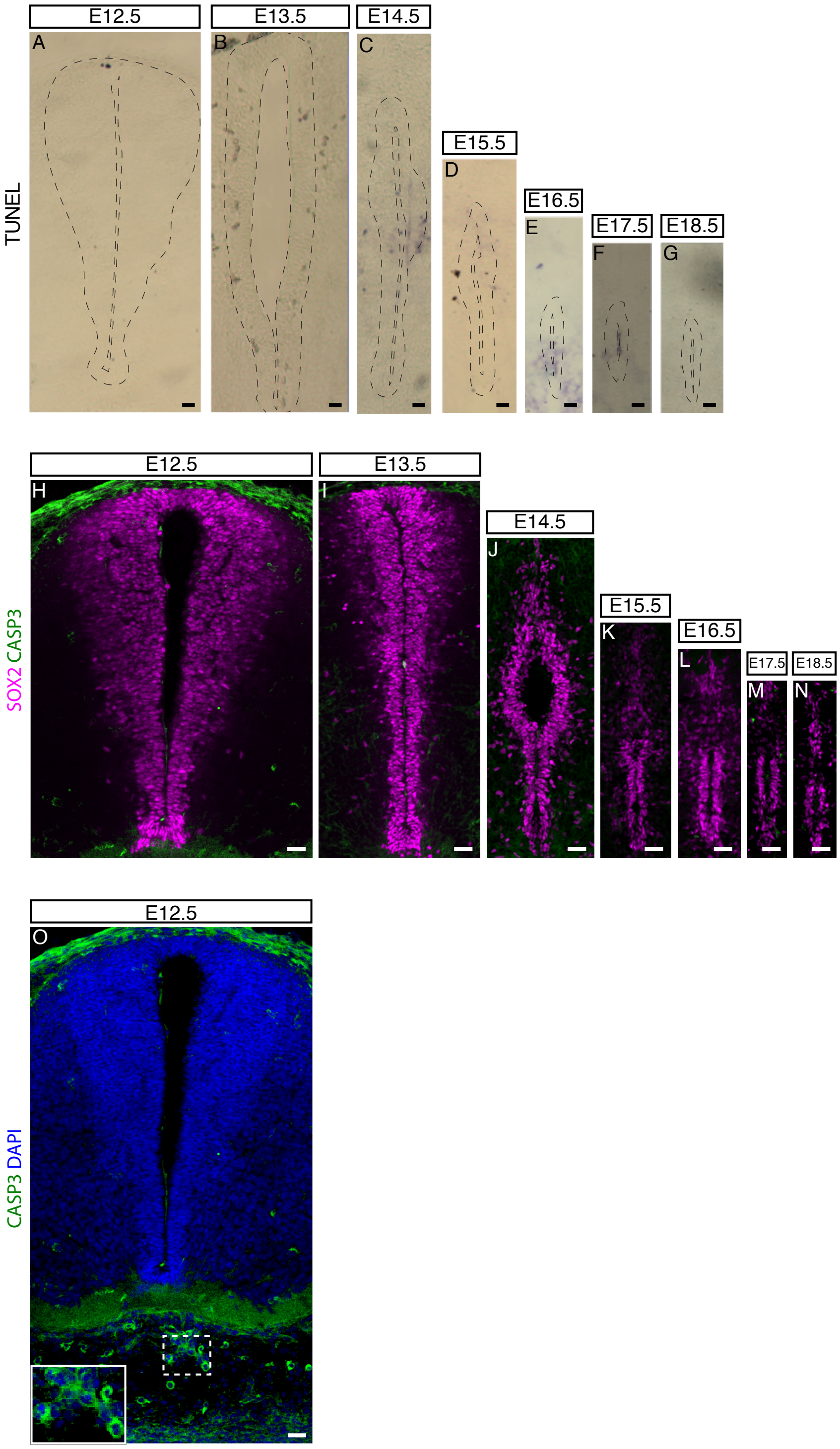


**Figure S1 Analysis of programmed cell death during mouse spinal cord development**
(A-G) TUNEL assay and (H-O) immunofluorescence for CASPASE-3 (green) was used to detect apoptotic cells at the indicated stages (9 sections, n = 3 embryos for each stage and each assay). White dashed box shows examples of CASPASE-3-positive cells found in the subarachnoid space at E12.5. Nuclei are stained with DAPI (blue). Progenitor cells within the ventricular layer are labelled with SOX2 (magenta). All scale bars: 20 μm.

### Supplementary tables

**Table S1** List of primary and secondary antibodies

Rabbit anti-ARX (1:1000, gift from Dr J. Chelly, IGBMC, Paris, France),
Mouse anti-BMPR1B (1:50, Santa Cruz Biotechnology sc-515886),
Mouse anti-CASPASE 3 (1:400, Abcam, ab13585),
Rabbit anti-FOXA2 (1:200, Abcam, ab108422),
Rabbit anti-FOXJ1 (1:200, Sigma, HPA005714),
Chicken anti-GFP (1:400, Abcam, ab13970),
Goat anti-GFP (1:400, Abcam, ab6673),
Mouse anti-PCNA (1:200, Merck Millipore, MAB424),
Rabbit anti-pH3 (1:400, CST, 9713),
Rabbit anti-pSMAD1/5 (1:50, CST, 9516),
Goat anti-SOX2 (1:200, Immune System, GT15098),
Rabbit anti-SOX2 (1:200-1:2000, Merck Millipore),
Mouse anti-TUJ1 (1:500, Covance Research Products Inc, MMS-435P),
Rabbit anti-TUJ1 (1:1000, Sigma, T2200).

All secondary antibodies were from Thermo Fisher Scientific Inc and diluted 1:500

Alexa 488 Donkey anti-rabbit IgG (H+L) Cat. # A21206;
Alexa 568 Donkey anti-rabbit IgG (H+L) Cat. # A10042;
Alexa 568 Goat anti-rabbit IgG (H+L) Cat. # A11011;
Alexa 594 Donkey anti-goat IgG (H+L) Cat. # A11058;
Alexa 594 Donkey anti-rabbit IgG (H+L) Cat. # A21207;
Alexa 594 Goat anti-rabbit IgG (H+L) Cat. # A11012;
Alexa 647 Donkey anti-goat IgG (H+L) Cat. # A21447.
Alexa 647 Donkey anti-mouse IgG (H+L) Cat. # A31571.

**Table S2** Plasmids, restriction endonucleases and RNA polymerases

| Plasmid – Gene product | Restriction endonucleases | RNA Polymerases |
|---|---|---|
| pYX-ACS-mFoxJ1 (full-length) | Anti-sense: SalI (Roche, Cat. # 10567663001)<br>Sense: Not1 (Roche, Cat. # 11014714001) | T3 (Roche, Cat. # 11031171001)<br>T7 (Roche, Cat. # 10881775001) |
| pCIG-mPtch1 (full-length) | SpeI (Roche, Cat. # 11008943001) | T3 |
| pBluescript II SK-mShh (insert size: 640 bp) | Anti-sense: HindIII<br>Sense: Not1 | T3<br>T7 |

**Tables S3-S6** are provided separately as Excel spreadsheets.
